# Resting-state functional connectivity relates to interindividual variations in positive memory

**DOI:** 10.1101/2021.06.20.448902

**Authors:** Ayako Isato, Tetsuya Suhara, Makiko Yamada

**Author notes:** Corresponding Author: Makiko Yamada, Institute of Quantum Life Science, National Institutes for Quantum and Radiological Science and Technology, Chiba, 263-8555, Japan. Declarations of interest: none.

## Abstract

Individual differences in positive memory recollection are of interest in mental health, as positive memories can help protect people against stress and depression. However, it is unclear how individual differences in positive memory recollection are reflected in brain activity in the resting state. Here, we investigate the resting-state functional connectivity (FC) associated with interindividual variations in positive memory by employing cluster-level inferences based on randomization/permutation region of interest (ROI)-to-ROI analyses. We identified a cluster of FCs that was positively associated with positive memory performance, including the frontal operculum, central operculum, parietal operculum, Heschl’s gyrus, and planum temporale. The current results suggest that positive memory is innervated by frontotemporal network connectivity, which may have implications for future investigations of vulnerability to stress and depression.

---

Individual differences in positive memory are of key interest in mental health as the recollection of positive memories can help people to cope with stress and enhance their wellbeing. As recalling positive past events can bring back pleasant feelings and neural activity of reward-processing, positive memory can protect individuals from stress and promote wellbeing. It has been shown that cortisol rise under stress exposure was mitigated in people who recalled positive, but not neutral, memories [1]. In older adults, positive memories contribute to their emotion regulation and wellbeing [2]. The severity of depressive symptoms is associated with reduced positive memory bias, in contrast to negative memory [3]. It has been shown that both currently depressed and remitted individuals had difficulty in recalling positive memories [4]. A recent study further revealed that recalling positive, but not negative memories, was predictive of a depressive mood and depressive vulnerability [5]. Thus, the ability to recall positive memories may be an important factor for mental health resilience.

In the past decades, a number of functional magnetic resonance imaging (fMRI) studies have revealed task-induced brain activity underlying positive memory, such as the cingulate gyrus and the bilateral frontal and parietal cortices [6]. Furthermore, positive memory performance was found to be related to post-encoding amygdala resting-state functional connectivity (FC), which increases within frontal regions [7]. Regarding depression, a current hypothesis suggests that positive memory reduction reflects disrupted communication between the mesolimbic dopamine pathway and medial temporal lobe memory systems during encoding [8]. However, it remains unclear which brain regions or networks are associated with interindividual variations in positive memory recollection. Brain activity in the resting state is known to mirror some task-induced brain activity [9] and can predict behavioral performance [10]. Considering the strong association between resting-state and task-induced brain activity, it is plausible to investigate the resting-state brain activity associated with positive memory.

Here, in order to elucidate the resting-state FC associated with individual differences in positive memory, we used spatial pairwise clustering (SPC) [11] which estimates the statistical significance level based on randomization/permutation ROI-to-ROI analyses, yielding less conservative cluster-based statistics compared to Bonferroni correction, the false discovery rate procedure, and extreme statistics.

Twenty-five right-handed healthy volunteers (22 men and 3 women) aged between 20 and 67 years (mean = 32.04, *SD* = 13.2) participated in this study. All participants provided written informed consent before participating in the study, which was approved by the Ethics Committee of the National Institute of Radiological Sciences and conducted in accordance with the ethical standards set forth in the 1964 Declaration of Helsinki and its later amendments. There are no conflicts of interest to declare.

In total, 102 pictures were taken from the International Affective Picture System [12]. These included: 34 positive pictures (e.g., delicious food, cute animals, sea landscape, baby, and sports), 34 negative pictures (e.g., gun, threatening animals, graves, and injured persons), and 34 neutral pictures (e.g., umbrella, marble pattern, fungus, and sober face). Of these, 51 pictures were presented in the incidental encoding task, and the same pictures together with the unlearned 51 pictures were presented in the surprise recognition test.

Participants underwent the incidental encoding task, where they did not explicitly intend to learn, followed by the surprise recognition test after 20 minutes. In the incidental encoding task, following the presentation of a fixation cross, a picture was presented for 750 ms (Figure 1A). Participants were asked to judge whether the picture was taken outside, inside, or unknown, by pressing response keys. A total of 51 pictures were presented in a pseudorandom order. After a 20 min intermission during which participants performed unrelated cognitive tasks, participants were asked to complete the recognition test. In this test, in which a picture was presented for 1500 ms (Figure 1B), all 102 pictures were presented in a pseudorandom order. Participants first judged whether they saw the picture in the incidental encoding task (“Old”) or not (“New”) by pressing response keys, and then rated their confidence level in their judgment using a 4-point Likert scale, ranging from “Not confident at all” to “Very confident”. After completing the surprise recognition test, participants performed the valence and arousal rating task, where each picture was presented for 1500 ms after the fixation cross (Figure 1C). They were asked to rate the valence and arousal of all the pictures with two 9-point Likert scales from “Very negative” to “Very positive” and from “Very calming” to “Very arousing”, respectively, using the Self-Assessment Manikin [13].

**Figure 1.**
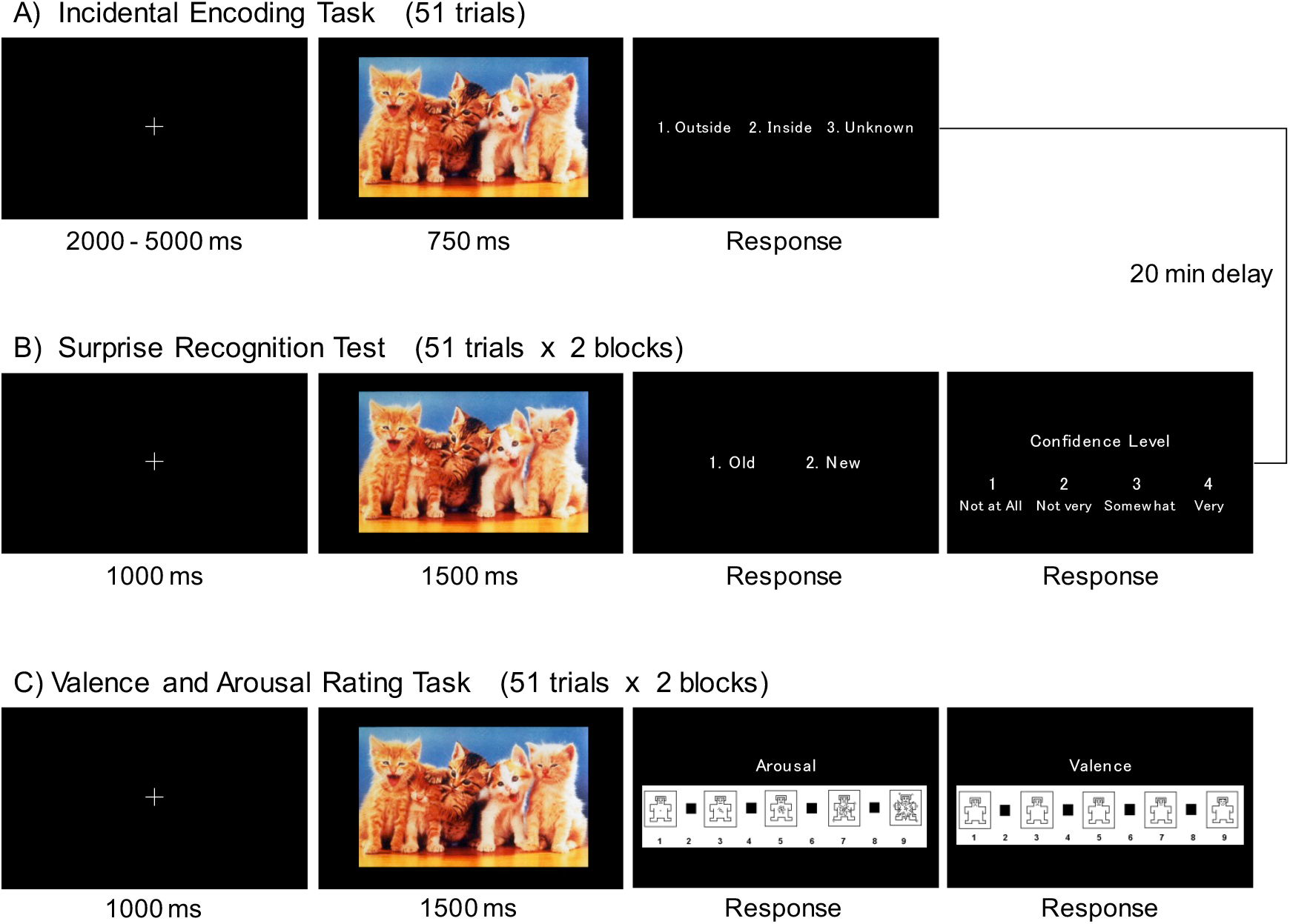
Trial structures of each task. (A) Incidental encoding task. Participants were asked to judge whether a picture was taken outside, inside, or unknown by pressing response keys. Fifty-one pictures including neutral, positive, and negative pictures were presented in pseudorandom order. (B) Surprise recognition test. The 51 pictures from the encoding task were presented again, along with 51 new neutral, positive, and negative pictures. Participants first judged whether they saw the picture in the incidental encoding task (“Old”) or not (“New”) by pressing response keys, and then rated their confidence level in their judgment using a 4-point Likert scale, ranging from “Not confident at all” to “Very confident”. (C) Valance and arousal rating task. Participants were asked to rate the valence and arousal of all the pictures with two 9-point Likert scales using the Self-Assessment Manikin (Bradley & Lang, 1994).

The responses obtained in the recognition test were divided into hit, miss, false alarm, and correct rejection. There were eight confidence levels in participants’ responses (from “New” and “Very confident” to “Old” and “Very confident”), and a borderline of these confidence levels was set as a criterion for the division of responses. For example, the response of “Old” and “Very confident” was categorized as “hit” if given for a picture that was presented in the incidental encoding task, but was categorized as “false alarm” if given for a picture that was not presented in the incidental learning task. The other response to a picture that was presented in the incidental learning task was categorized as “miss”, and the other response to a picture that was not presented in the incidental learning task was categorized as “correct rejection” when that response was set as a borderline. Based on the signal detection theory, the receiver operating characteristics (ROC) curve of each participant was generated by plotting cumulative hit rates on the y-axis and false alarm rates on the x-axis for every confidence level [14]. The area under the curve (AUC) of the ROC curve was used as an index of the recognition test performance. Separate one-way repeated measures ANOVAs with Bonferroni *post hoc* correction (positive vs. negative vs. neutral) were applied to examine the differences between emotions in the performance of the recognition test (AUC) and the ratings of valence and arousal.

All participants underwent a resting-state fMRI scan. A Siemens Verio MRI system (3T) was used to obtain T2*-weighted echo-planar imaging (EPI; repetition time = 2,000 ms, echo time = 25 ms, slice number = 33, thickness = 3.8 mm, matrix = 64 × 64, slice gap of 14 out of 25 participants [11 out of 25 participants] = 0 [0.456] mm, 204 volumes, interleaved acquisition) and structural T1 images (repetition time = 2,300 ms, echo time = 1.95 ms, voxel size = 1 × 0.488 × 0.488, slice number = 176) with a dedicated 32-channel head coil. During resting-state fMRI runs, participants were instructed to lie as still as possible for approximately 7 min and to maintain gaze on the fixation cross without thinking of anything.

To process fMRI data, we used the Functional Connectivity (CONN) toolbox (https://web.conn-toolbox.org/), which was implemented in Statistical Parametric Mapping software version 12 (https://www.fil.ion.ucl.ac.uk/spm/software/spm12/). Realignment, slice-timing correction, normalization, and smoothing with Gaussian filter kernel (full width at half maximum = 6 mm) were conducted. Before pre-processing, the first four volumes were left out. Conservative functional outlier detection settings were used during the Artifact Detection Tools (ART)-based identification of outlier scans for scrubbing (global signal z-value threshold of 3, subject-motion threshold of 0.5 mm). WM, CSF, and realignment parameters were entered as confounds in a first-level analysis, and a band-pass filter (0.009–0.1 Hz) was applied.

ROI-to-ROI analysis was conducted to examine which FC is implicated in interindividual variations in positive emotional memory using the CONN toolbox. We predefined 105 ROIs in the CONN toolbox based on the Harvard–Oxford cortical and subcortical structural atlases (https://neurovault.org/collections/262/). The brainstem and the cerebellum were excluded. We selected statistically significant ROI-to-ROI connections that correlate with the performance of the recognition tests by cluster-level inferences based on spatial pairwise clustering statistics (SPC) [11] with default settings in CONN. The T-statistics of the entire ROI-to-ROI matrix were estimated using a general linear model. Age, gender, and the performance in the recognition test (AUC) of neutral and negative pictures were defined as the confounding covariates, and that of positive pictures as the covariates of interest to examine ROI-to-ROI connections that relate to interindividual variations in positive emotional memory. ROIs in this matrix were sorted automatically by a hierarchical clustering procedure (optimal leaf ordering for hierarchical clustering, [15]) based on ROI-to-ROI anatomical proximity or functional similarity metrics. Thereafter, this statistical parametric map was thresholded using a significance level of *p* < .01 (uncorrected). The resulting suprathreshold areas define a series of non-overlapping clusters. The cluster-level FDR-corrected *p* < .05 was applied.

One-way repeated measures ANOVA for the AUC revealed a significant main effect of emotion (*F*(2,48) = 5.023, *p* = 0.010, η_p_^2^ = .173, Neutral: *M* = 0.90, *SD* = 0.08, Positive: *M* = 0.87, *SD* = 0.09, Negative: *M* = 0.92, *SD* = 0.05). The Bonferroni *post hoc* test indicated a significant difference between positive and negative recognition (*p* = 0.006), but no other differences were found.

One-way repeated measures ANOVA for valence ratings revealed a significant main effect of emotion (*F*(2,35.928) = 209.034, *p* < 0.001, η_p_^2^ = .897, Neutral: *M* = 5.05, *SD* = 0.67, Positive: *M* = 6.47, *SD* = 0.86, Negative: *M* = 2.80, *SD* = 0.70). The Bonferroni *post hoc* test showed that the valence of positive pictures was rated significantly higher than that of negative and neutral pictures, and the valence of neutral pictures was rated significantly higher than that of negative pictures (all *p* < 0.001).

One-way repeated measures ANOVA for arousal ratings revealed a significant main effect of emotion (*F*(2,36.075) = 66.152, *p* < 0.001, η_p_^2^ = .734, Neutral: *M* = 1.62, *SD* = 0.76, Positive: *M* = 2.06, *SD* = 1.24, Negative: *M* = 4.46, *SD* = 1.73). The Bonferroni *post hoc* test showed that the arousal of negative pictures was rated significantly higher than that of positive and neutral pictures (all *p* < 0.001), and the arousal rating of positive pictures was higher than neutral pictures at a marginally significant level (*p* = 0.055). Thus, based on the behavioral analyses, we confirmed that the three sets of pictures were successfully perceived as positive, negative, and neutral, as defined.

To extract the FCs specific to positive memory, we performed resting-state FC analysis using the SPC by including age, gender, and AUCs of neutral and negative pictures as covariates of no interest and the AUC of positive pictures as a covariate of interest. The SPC analysis revealed a total of 52 clusters of ROIs at the t-statistic threshold (*p* < .01), and one cluster of FCs was identified by cluster-level FDR-corrected p-values (*p* < .05). The FCs included the right and left frontal operculum cortex, right and left central opercular cortex, right and left planum temporale, right and left Heschl’s gyrus, and left parietal operculum cortex (Table 1 and Figure 2).

**Table 1.**
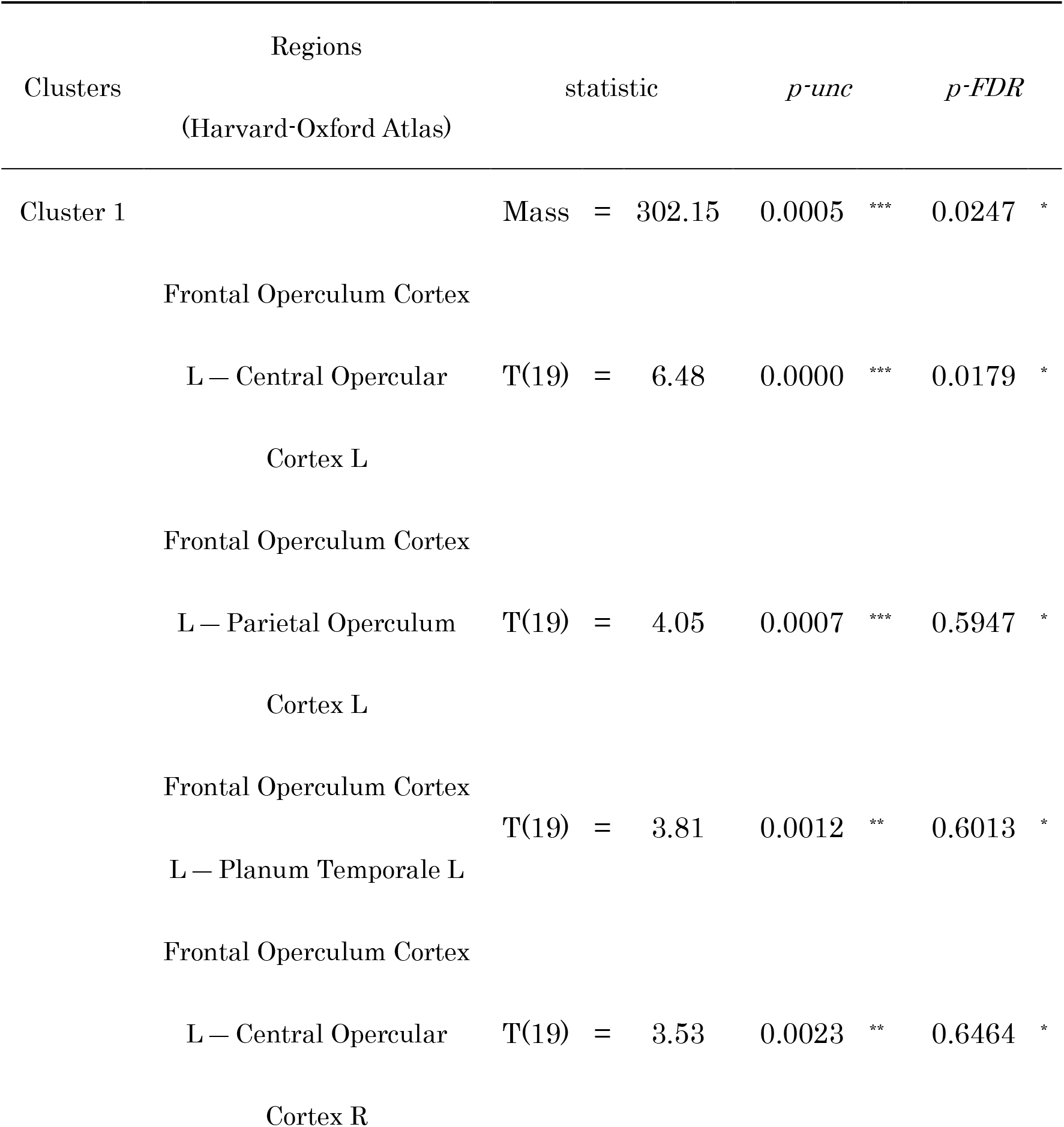
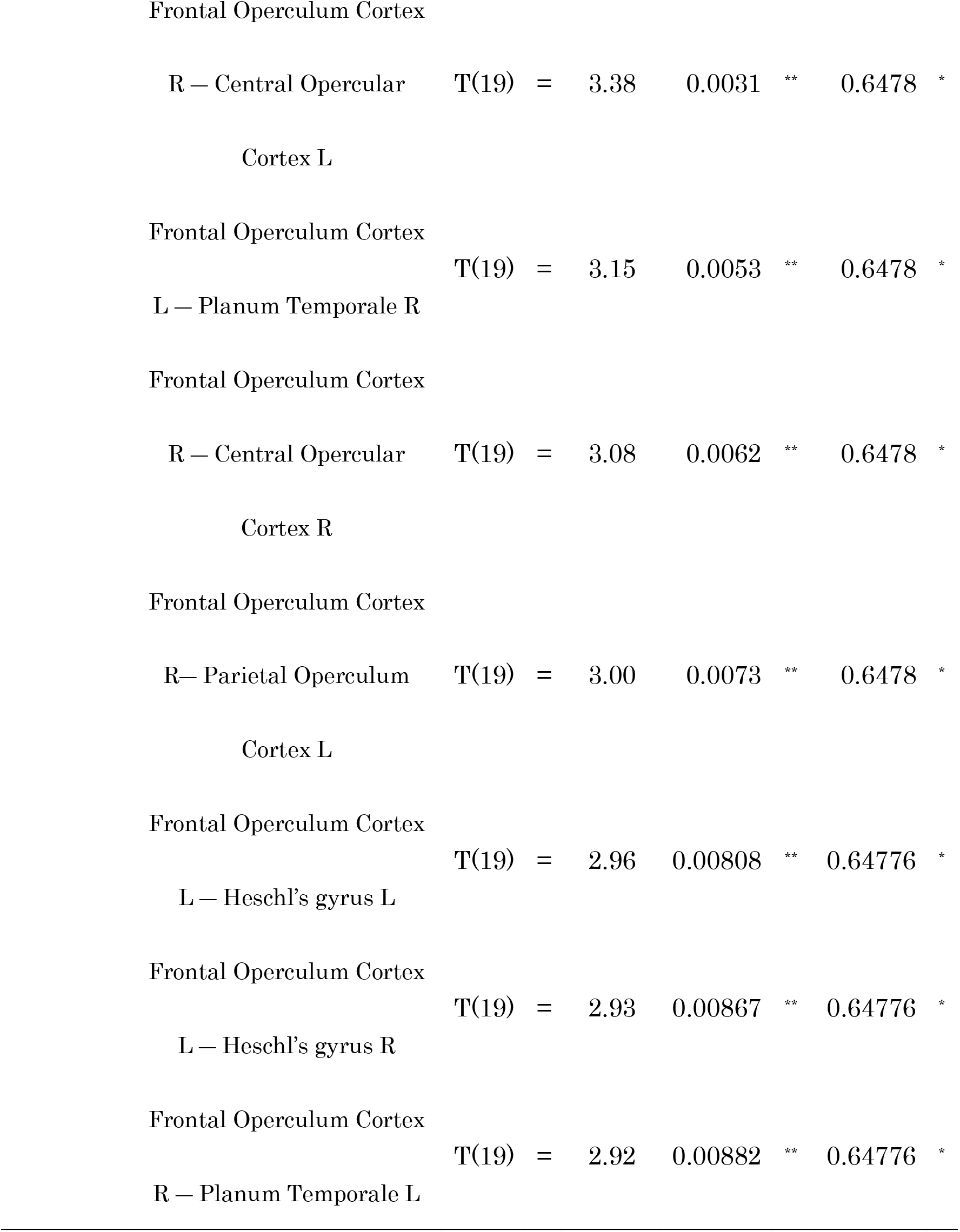
A cluster of ROI-to-ROI connections showed a significant positive correlation with the performance of the recognition test of positive pictures.

**Figure 2.**
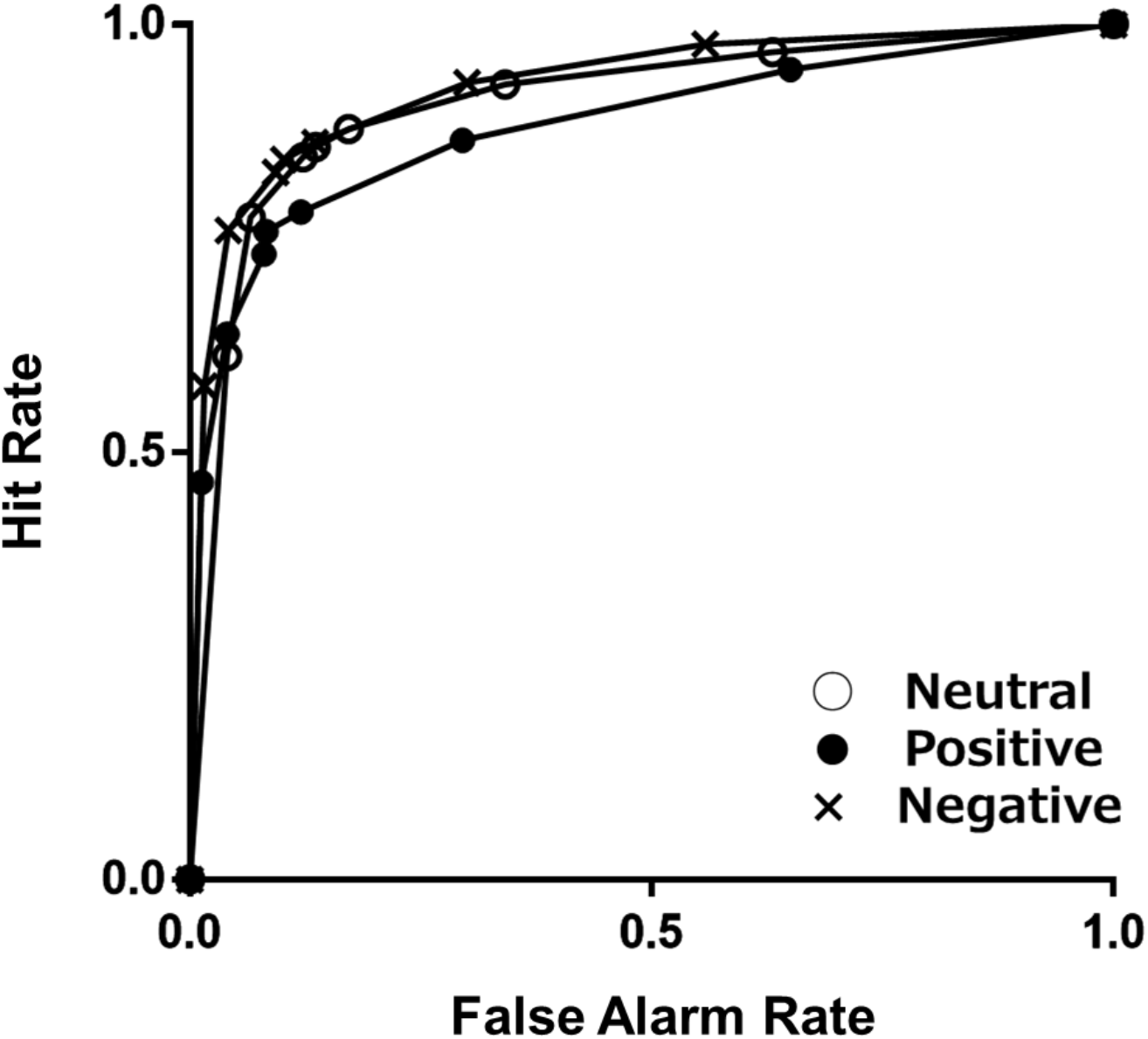
ROC curves relating the recognition hit rate to the false alarm rate for neutral, positive, and negative pictures.

**Figure 3.**
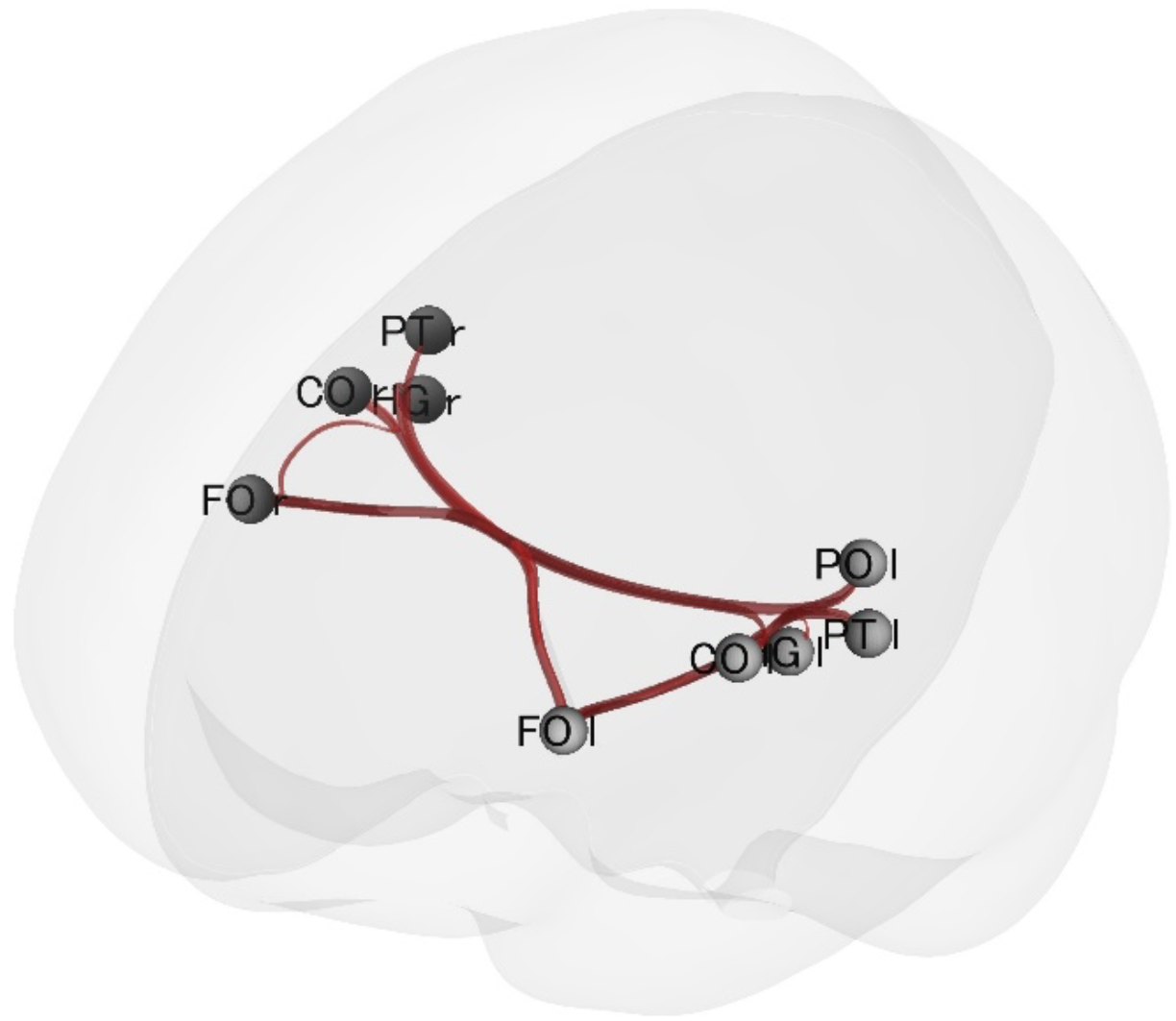
Clusters of FCs that show significant positive correlations with the recognition ability of positive pictures. The thickness of the lines shows the magnitude of the correlation.

The identified frontotemporal network connectivity was positively correlated with the recollection of positive memories. Previous task-based fMRI studies revealed that frontotemporal regions were associated with the processing of positive and auditory stimuli, such as the processing of pleasant music [16], multimodal integration such as emotional and auditory information in musical performance [17], and positive emotions, especially joy [18]. Considering that positive memory is likely to be enhanced in verbal or conceptual information [19], the language functions of the frontotemporal networks may play an important role in positive memory recollection.

In general, positive emotions play a protective role in terms of mental health. Recalling positive memories may be a protective mechanism that facilitates physiological and emotional stress recovery [1], and both depressed and at-risk adolescents show positive autobiographical memory deficits [20].

To support the above view, it has been shown that a negative memory bias in patients with depression was replaced by a positive memory bias after electroconvulsive therapy (ECT) [21]. Moreover, positive memory enhancement training, which prompts participants to recall vivid and detailed positive memory, had the effect of improving mood in individuals with major depressive disorder [22]. Thus, the current finding that the frontotemporal network connectivity is associated with positive memory recollection suggests that this network may be a possible target for functional connectivity neurofeedback training [23], aiming to enhance positive memory to safeguard against stress and depression.

The major limitation of the current study is the imbalance of gender ratio. Although gender was included as a covariate, the gender differences in emotion processing are widely known. Thus, future studies may want to clarify whether FCs found here vary by gender.

In summary, the current study revealed that the frontotemporal network connectivity was exclusively associated with the recollection of positive memories in healthy individuals. Future studies are required to clarify the role of this network in positive memory impairments in patients with depression, which may influence both depressive vulnerability and symptom severity.

